# Phylogeographic patterns of *Escherichia coli* isolated from white-tailed deer, *Odocoileus virginianus*, during necropsy in a farmed deer surveillance program

**DOI:** 10.1101/2024.11.21.624738

**Authors:** Morgan C. Metrailer, An-Chi Cheng, Andrew P. Bluhm, Treenate Jiranantasak, Michael H. Norris, Austin Surphlis, Kuttichantran Subramaniam, Kwangcheol Casey Jeong, Samantha M. Wisely, Juan Campos Krauer, Jason K. Blackburn

## Abstract

White-tailed deer (*Odocoileus virginianus*) (WTD) farms are distributed throughout Florida and number nearly 400. The Cervidae Health Research Initiative (CHeRI) is an initiative that investigates disease and health of farmed cervids statewide. Within the Florida industry, Epizootic Hemorrhagic disease virus (EHDV) and Bluetongue virus (BTV) are major causes of disease and associated economic losses. Often, deer survive EHDV or BTV infections but succumb to secondary bacterial infections, including *Escherichia coli. E. coli* can be opportunistic with severe illness or death occurring in hosts weakened from viral infection. We investigated presumptive *E. coli* strains isolated from WTD during necropsy investigations. Genomic DNA was extracted and whole genome sequenced from 61 suspect *E. coli* isolates. Exploratory spatial data analysis (ESDA) of the distribution of *E. coli* phylotypes, distribution of *E. coli*, and deer between ranches was performed. We examined phylogenetic relationships between isolates, categorized by ranch and year and compared to all reported *E. coli* isolates in Florida from EnteroBase. One strain was *Enterobacter hormaechei* and the other 60 were *E. coli*. Two *E. coli* strains were toxigenic. Deer isolates represented 7 phylogroups with B1 being the most prevalent (45/60) and geographically widespread (14/16 counties reporting *E. coli*). Phylogroup A was the second most prevalent phylogroup. In at least two instances two phylogroups were present within a single animal. We found that deer isolates spanned many of known phylotypes in Florida from animals and humans. Diversity patterns suggest deer are infected locally with most animals on ranches having high genomic similarity within phylogroups.

*Sequence data from this article have been deposited in the National Center for Biotechnology Information (NCBI) database under BioProject PRJNA1183872.

## INTRODUCTION

White-tailed deer (*Odocoileus virginianus*) (WTD) are economically important species due to their widespread abundance and their prominence in the U.S. hunting industry [1]. Within Florida, WTD ranches number nearly 400 – with calls of concern by ranchers and owners of morbidity and mortality due to disease. The Cervidae Health Research Initiative (CHeRI) is a multi-disciplinary effort enacted by University of Florida IFAS to monitor and diagnose hemorrhagic disease (HD), a leading cause of death in Florida captive cervids (CHeRI; University of Florida, Gainesville, FL, USA). Epizootic hemorrhagic disease virus (EHDV) and blue tongue virus (BTV) are the primary causative agents of HD in Florida WTD [2]. However, infections caused by bacterial pathogens, typically *Escherichia coli* and *Clostridium* spp., were among the most prevalent in young fawns (between 2 – 6 weeks old) [3]. These pathogens can also be attributed to secondary infections in adult WTD that have been weakened by active or recently passed viral infections.

*Escherichia coli* is a highly studied Gram-negative bacterium that occupies a diverse range of niches [4,5]. Classically, *E. coli* strains have been placed into seven genetic phylogroups (A, B1, B2, C, D, E, and F) [6]; with pathogenic strains primarily falling into groups B2 and D [7–9]. Phylogroups A and B1 were more commonly commensal but also have shown to have greater antimicrobial resistance (AMR) [10–12]. While the commensal nature of most *E. coli* strains can prove challenging in epidemiological traceback or geographic placement, there are some established patterns associated with specific phylogroups [13,14]. Often, certain hosts species or isolation sources (i.e., the environment) are associated with specific phylogroups and the identification of uncommon phylogroups within a host may indicate potential co-mingling or higher exposure to human modified environments [4]. Classification of *E. coli* strains into these broad scale (lower genetic resolution) phylogroups can also provide initial indications of potential pathogenicity or risk of AMR. Intermingling of strains within these phylogroups pose a risk as pathogenic strains may attain AMR genes (or vice versa) through horizontal gene transfer [8,10].

In 2000, a triplex polymerase chain reaction (PCR) assay targeting genes *chuA* and *yjaA* and DNA fragment TSPE4.C2 was developed to rapidly distinguish groups A, B1, B2, and D [15]. Expansion of multi-locus sequencing typing (MLST) of *E. coli* house-keeping genes clarified strain classification into phylogroups A, B1, B2, and D and led to the classification of phylogroups E, F, and C [16]. In 2013, Clermont et al. improved the original triplex primers to limit the impact of single nucleotide polymorphisms (SNPs) and to exclude *Escherichia* cryptic clades [6]. The assay was also expanded to a quadplex PCR, with the addition of primers targeting the *arpA* gene to discriminate group F. Following the quadplex, additional primers or the use of MLST could be used to distinguish group C and group E [6]. In 2019, primers were produced to classify strains in the newly identified group G [17]. As whole genome sequencing became more accessible, these *in vivo* methods were translated into *in silico* methods [18]. A web-based classification system was produced, EzClermont, for rapid phylotyping of *E. coli* genomes [19]. This rapid typing method allows for easier placement of WGS *E. coli* genomes and links current modern genomes to traditionally typed isolates (studies where WGS was not available).

A core genome MLST (cgMLST) utilizes more of the genome than previous described methods and has greater variability. The current *E. coli* specific cgMLST utilized by Enterobase was seeded from *E. coli* K-12 (Accession number: NC_000913.3) and identified 2,513 loci within the core genome [20]. Within each loci there is a distinct DNA sequence, classified as an allele type. The total cgMLST profile is then used to assign a strain, or isolate, to a sequence type (ST) based on these core genes. Agglomerative clustering methods have been used to investigate genetic relationships between strains where each isolate begins it’s own cluster. A linkage function is then used to determine how individual ‘clusters’ may pair. Single-linkage clustering takes a single element from each cluster that have the lowest pairwise distance and merges them (nearest neighbor clustering) [21]. Similarly, minimum spanning trees (MSTs) utilize these agglomerative (bottom-up) clustering methods to form trees in which the minimum distance between individual nodes are linked until all nodes fall within the tree. A multi-level (hierarchical) clustering scheme was integrated into Enterobase [21]. This algorithm first produces minimum spanning trees from the cgMLST schema, accounting for any missing locis. It then re-assigns nodes (individual genomes) to the lowest possible cluster designation if they have equal genetic distances to multiple clusters in the tree [21].

This study investigates the spatial and phylogenetic patterns of WTD mortalities and their associated diseases through cooperative diagnostic programs implemented by UF IFAS CHeRI. While viral diseases, such as EHDV and BTV, are established and endemic in these populations [2], the burden of disease caused by commonly commensal bacteria, like *E. coli*, can be elusive. The objective of this study was to 1) identify the spatial patterns of white-tailed deer mortalities and their associated *E. coli* isolates, 2) determine the role of *E. coli* as a potential secondary infection and the distribution of pathogenic *E. coli* or neighboring *Enterobacter* species and, 3) investigate *E. coli* phylogenetics at multiple genetic and spatial scales.

## METHODS

The total number of licensed white-tailed deer ranches in Florida, as of 2023, were identified and geocoded. White-tailed deer ranch density per county was calculated and choropleth mapped in QGIS. From 2020 – 2022, the proportion of these ranches that participated in CHeRI project was calculated and choropleth mapped. Dead farmed white-tailed deer from participating CHeRI farms across Florida were sampled to test for the presence of bluetongue virus (BTV), epizootic hemorrhagic disease virus (EHDV) and *Escherichia coli* (*E. coli*). Samples from different organ tissue (kidney, heart, liver, spleen) and whole blood were collected by the local deer farmers or by CHeRI staff and transported to CHeRI laboratories on ice or cold packs. The number of deer mortalities per county was calculated and the density was choropleth mapped.

Suspected *E. coli* colonies were cultured and isolated from organ tissue and saved into glycerol stocks. Frozen glycerol stocks were inoculated into brain-heart infusion (BHI) broth (BD Difco™ Bacto™ or BD Difco™ 237500) and incubated at 37°C with shaking (220 RPM) overnight. Genomic DNA was extracted using the Wizard Genomic DNA purification kit (Promega Corporation, Madison, WI, USA) and quantified using the Qubit 1X dsDNA high sensitivity kit (Invitrogen, Waltham, MA, USA). The smoothed distribution of recovered *E. coli* isolates from each ranch was produced using a kernel density estimation with the local neighborhood bandwidth being determined using the h_opt_ formula [22]. The resulting distribution was mapping over the Florida county boundaries.

Recent and/or active infections with Epizootic Hemorrhagic Disease virus (EHDV) or Bluetongue virus (BTV) were identified with a reverse transcriptase (RT) qualitative PCR (RT-qPCR) assay [2,23]. From the WTD mortalities, splenic tissue was collected, and a sample was frozen at −80°C for downstream viral analyses. Total RNA was extracted using the Qiagen RNeasy Mini kit (Qiagen, Valencia, CA, USA) and screened using the Applied Biosystems VetMAX Plus One-Step RT-qPCR kit (Applied Biosystems, Invitrogen, Waltham, MA, USA). A histogram depicting the number of *E. coli* isolates recovered from WTD hosts with no identified co-infections (*E. coli* only) and with identified co-infections (with EHDV, with BTV, or with both EHDV/BTV) was produced using Microsoft Excel (Microsoft, Redmond, WA, USA). The co-infection status of recovered isolates was visualized as proportions (pie-charts) for each ranch and mapped.

Whole-genome sequencing of suspected *E. coli* isolates was performed using the Illumina NovaSeq platform following the construction of sequencing libraries using the Illumina DNA Prep kit (Illumina, San Diego, CA, USA). The quality of the resulting paired-reads was assessed using FASTQC [24]. Raw reads were uploaded to Enterobase for downstream phylogenetic analyses. Enterobase is a large online database consisting of more than 1 million bacterial genomes, with a majority of the strains being enteric bacterial species, *Salmonella, Escherichia*, and *Shigella*. Here, the raw sequencing reads were assembled, and the identification of virulence genes (VG) was performed using BlastFrost [25]. Specifically, the identification of Shiga toxin genes (*Stx1, Stx2*), enteroinvasive virulence genes (*pInv)*, heat stable endotoxin genes (ST), heat-labile endotoxin genes (LT), and adhesion genes (*eae*). Within Enterobase, the *E. coli* pathotype was determined based on the presence of these virulence genes. Separately, genome assembly of *E. coli* isolates was performed using a snakemake pipeline employing trimmomatic [26], to remove adapter sequences and low quality reads, and spades assembler [27], optimized with Unicycler [28]. Quality of assembled genomes was confirmed using QUAST against reference strain from *E. coli* K-12 (Accession number: NC_000913.3). Phylogroup classification of isolates was determined using the *in silico* EzClermont web-based tool [19]. If the QUAST assembly statistics and phylogroup classification was inconsistent with *E. coli*, genus and species was determined using the PubMLST Species Identifier online tool (https://pubmlst.org/bigsdb?db=pubmlst_rmlst_seqdef_kiosk). Ranches that had WTD mortalities containing potentially pathogenic isolates were identified and mapped.

A neighbor-joining phylogeny of the recovered *E. coli* isolates was produced in Enterobase using their hierarchical core genome multi-locus sequence typing (MLST) scheme (HierCC+cgMLST) [21]. The resulting tree was visualized with their assigned phylogroup and year of isolation using Interactive Tree of Life (iTOL). The spatial distribution and county level phylogroup diversity was mapped using QGIS. A phylogeny of all Florida isolates listed in Enterobase (n=596) was produced using the HierCC+cgMLST scheme. The tree was visualized in iTOL with branch colors representing phylogroup designation and two outer rings; host (or isolation source) and if the strain was a CHeRI isolate. A minimum spanning tree (MST) of all Florida isolates was also produced with the CHeRI isolates indicated in black. A nested view of the B1 phylogroup branch was incorporated and colored according to ranch and deer number.

## RESULTS

### CHeRI participation and sample recovery

Counties containing white-tailed deer (WTD) ranches are distributed throughout Florida, with central and north central Florida counties housing the greatest amount (Figure 1a). There is gapped coverage both at the country level (0 ranches reporting to CHeRI) or only partial coverage within counties (blue – purple counties) (Figure 1b). From 2020 – 2022 the number of white-tailed deer mortalities was greatest in the northern panhandle region (n=16) (Figure 1c). The counties reporting the most mortalities during this period were Calhoun and Gadsen (shown in black) (Figure 1c). From these, the smoothed KDE showed that the greatest number of recovered *E. coli* strains was also in this panhandle region, comparable to the number of mortalities (Figure 1d).

**Figure 1.**
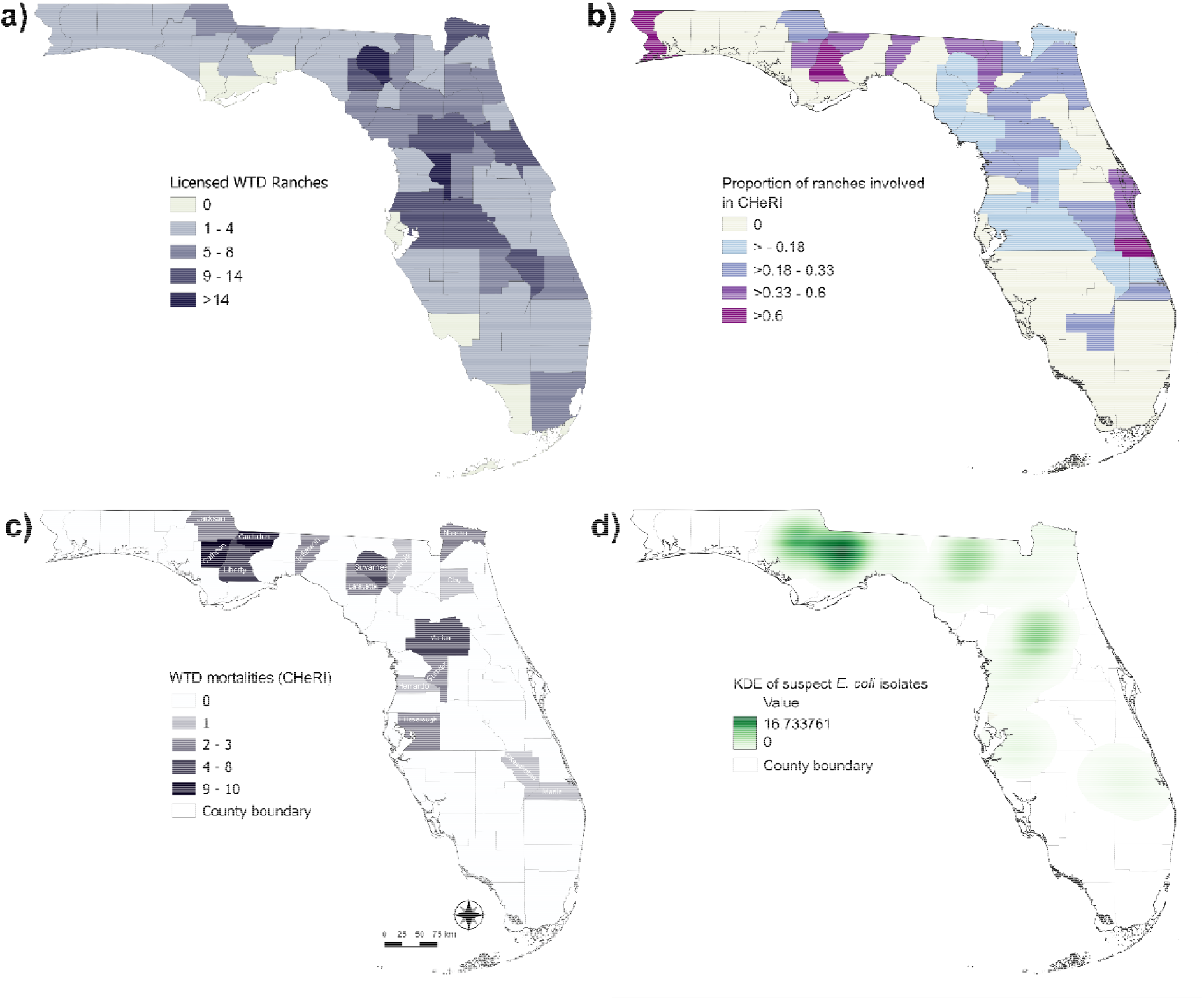
Spatial patterns of white-tailed deer (WTD) ranches and *E. coli* samples in current CHeRI project. Count of licensed white-tailed deer ranches per Florida counties (a). Proportion of licensed white-tailed deer ranches involved in the Florida CHeRI project to total WTD ranches per counties (b). Counties with no white-tailed deer ranches are shown in tan. Choropleth map of Florida counties with counts of white-tailed deer deaths submitted to CHeRI in 2023 (c). Kernal density estimation of suspected *E. coli* isolates from white tailed deer mortalities submitted to CHeRI in Florida (d). Bandwidth determined using the h_opt_ formula [22].

### Spatial distribution of co-infections and pathogenic isolates

A majority of *E. coli* isolates (n=35) were recovered from WTD mortalities that were negative for both EHDV and BTV (Figure 2a, gray). In the isolates that had co-infections, a subset was positive for EHDV (n=13; Figure 2a, light purple) or BTV (n=7; Figure 2a, purple). Five *E. coli* isolates were recovered from WTD mortalities that were positive for both EHDV and BTV (Figure 2a, dark purple). A majority of ranches, distributed throughout the study area, did not report deer mortalities with co-infections with EHDV and/or BTV (Figure 2a, gray pie charts). Co-infection status was not restricted within each ranch – WTD mortalities with no identified viral infections were present on ranches with mortalities positive for EHDV, BTV, and both EHDV/BTV. The distribution of these co-infection ranches was not geographically constrained; however, the greatest diversity of co-infection status was reported in the northern panhandle (Figure 2a). Ranches with potentially pathogenic isolates were also located in the panhandle region (Figure 2b). Two enterotoxigenic (ETEC) *E. coli* strains, both having heat stable endotoxin genes (ST), were identified at two ranches in separate counties (Figure 2b, red). A third ranch recovered a suspected *E. coli* strain that was later identified as *Enterobacter hormaechei* (Figure 2b, purple).

**Figure 2.**
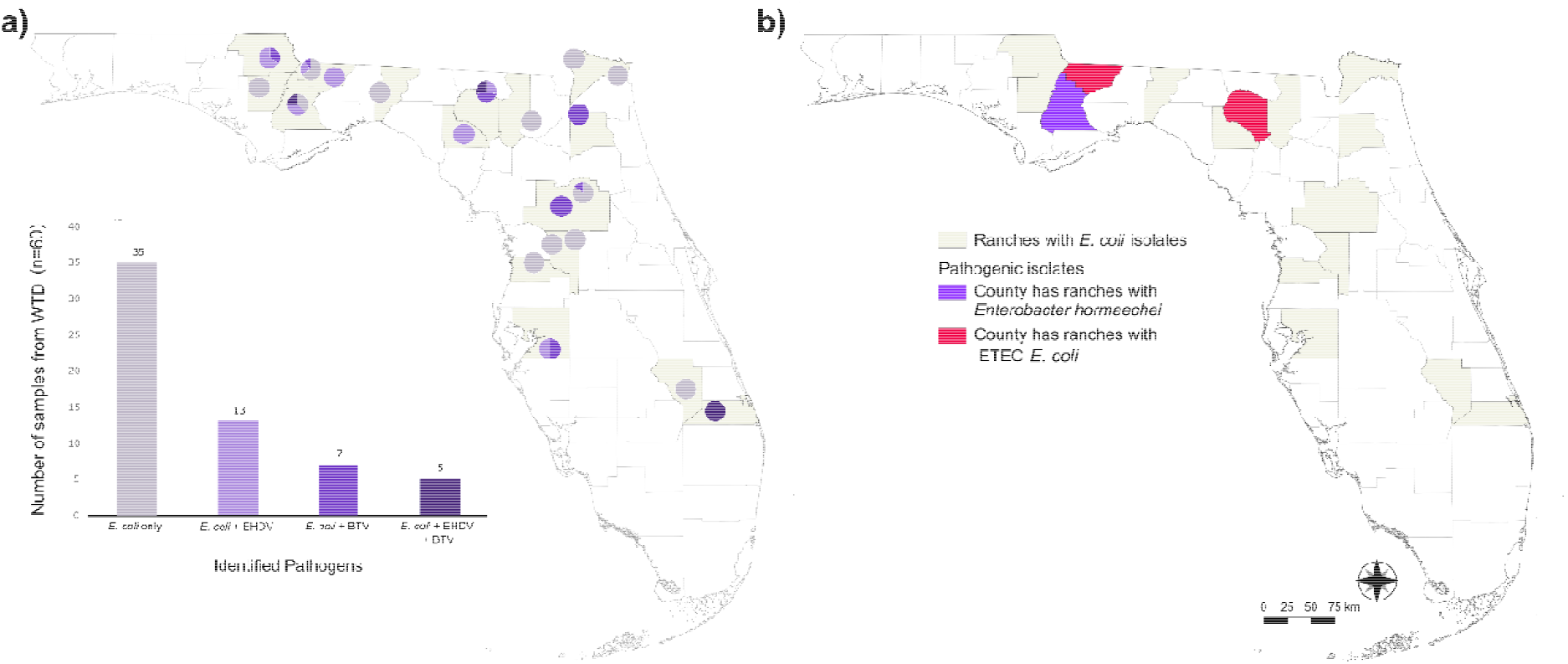
Viral co-infections and pathogenicity of *E. coli* isolates in 2023 Florida study. Distribution of Florida counties with pie charts depicting co-infection with Bluetongue Virus (BTV; purple), Epizootic Hemorrhagic Disease Virus (EHDV; light purple), or both viruses (dark purple) (a). Pie charts are mapped to approximate ranch location and dithered. *Escherichia coli* isolates from mortalities negative for BTV and EHDV are shown in gray. Florida counties with Enterotoxigenic (ETEC) *Escherichia coli* (red) and *Enterobacter hormaechei* (purple) isolates identified in this study period (b).

### Phylogenetics: spatial patterns and diversity

The most prevalent phylogroup in this study was Phylogroup B1 (Table 1, n=45; Figure 3, orange). This phylogroup was dispersed across the study area – identified in 17 ranches, spanning 14 counties (Table 1). Phylogroup A accounted for 7 strains, spanning 5 ranches in 4 counties (Table 1, n=45; Figure 3, pink). Phylogroups D (blue) and E (light green) were identified for 3 and 2 strains, respectively (Table 2, Figure 3). Phylogroups B2 (green), C (yellow), and F (red) were only identified in one isolate each (Table 2, Figure 3). Counties with the greatest phylogroup diversity were in the northern panhandle region of the study area. Isolates from the white-tailed deer (family: Cervidae) were closely related to *E. coli* isolates from Bovine species, Equine species, and humans (Figure 4). The most prevalent phylogroups identified in all Enterobase *E. coli* Florida isolates were B1 and B2 with phylogroups C, D, E, and F remained the rarest phylogroups in the phylogeny (Figure 4).

**Table 1.**
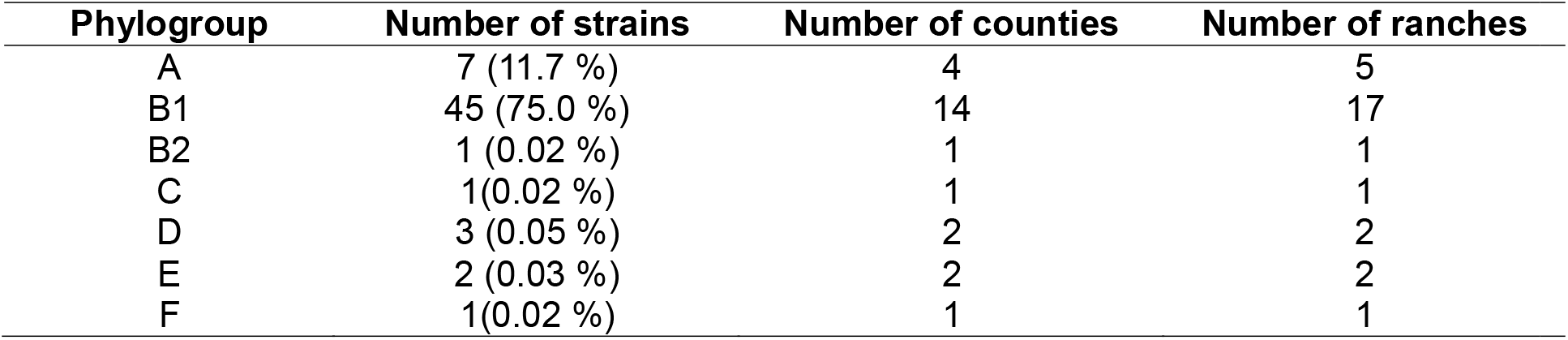
*Escherichia coli* phylogroups identified in this study. The number and percentage of strains within the phylogroup is listed. The number of ranches (n=20) and counties (n=16) that identified these phylogroups is reported. Total number of recovered *E. coli* isolates in this study was 60. One additional isolate was recovered but later identified as *Enterobacter hormaechei*.

**Figure 3.**
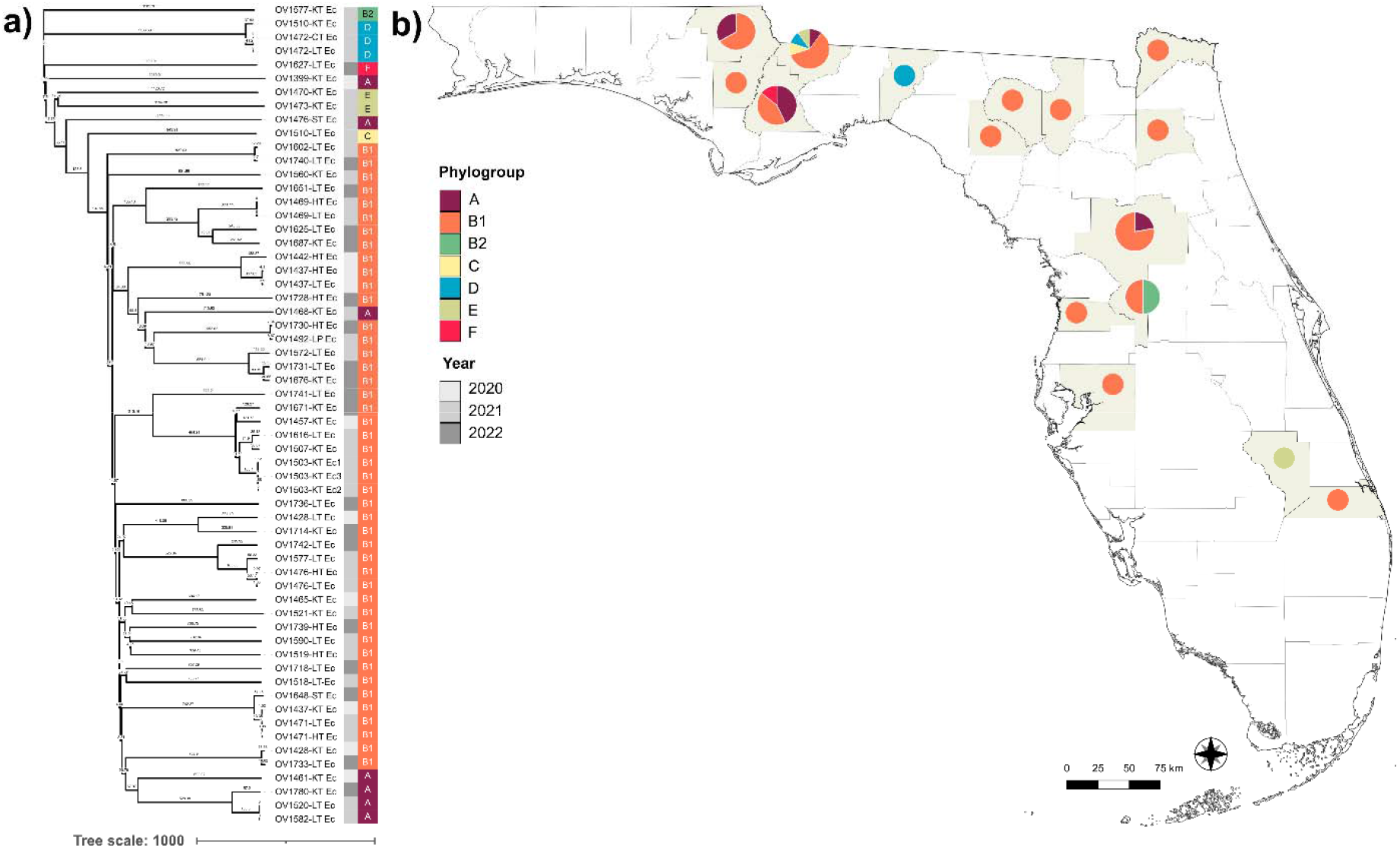
*Escherichia coli* phylogeny of CHeRI isolates (a) with the spatial distribution of *E. coli* phylogroups diversity per county (b). Neighbor-joining phylogeny was produced using the Enterobase HierCC+cgMLST algorithm and visualized in iTOL. Year of isolate recovery from WTD mortality is shown in gray (2020-2022).

**Figure 4.**
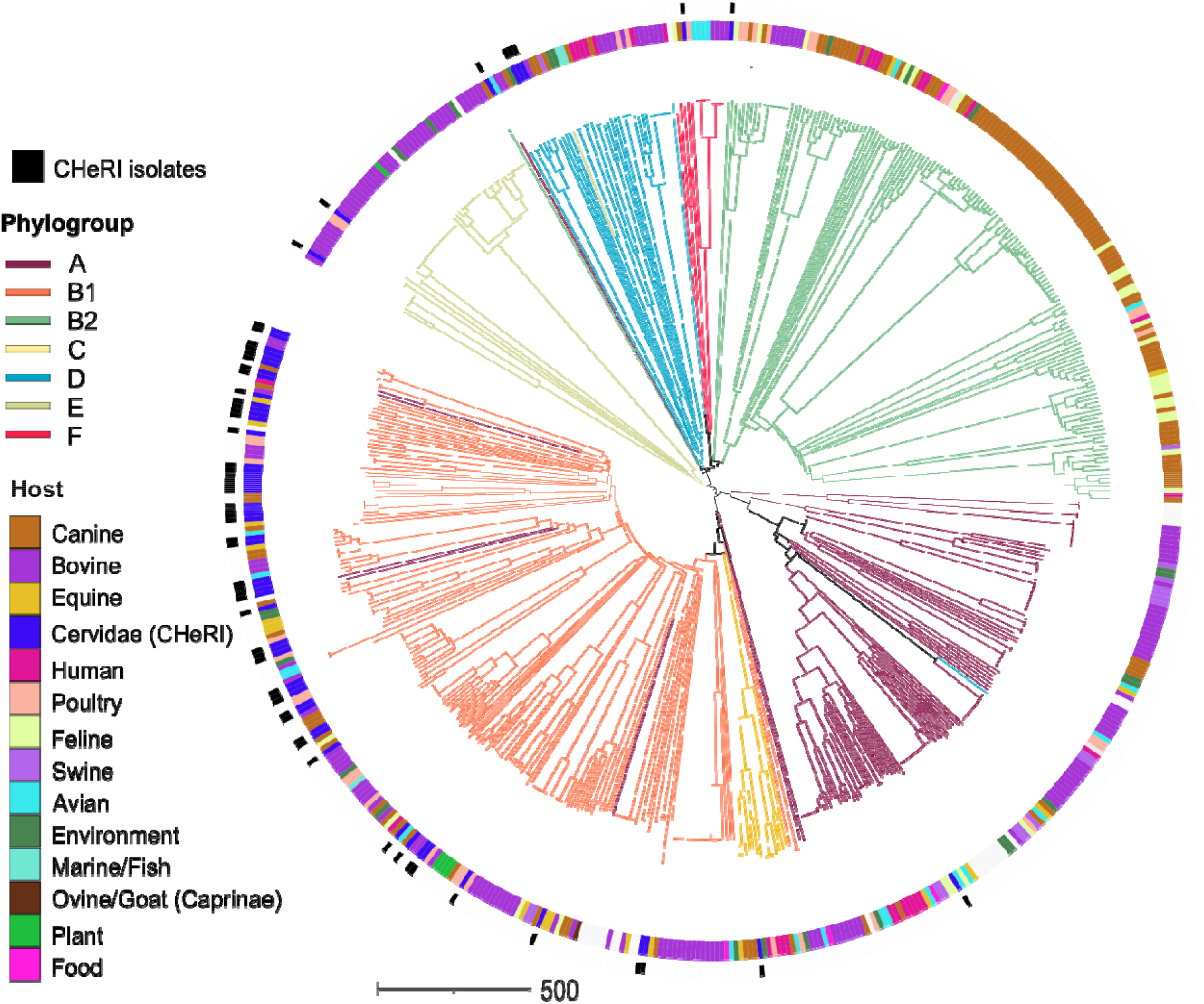
Core-genome MLST (cgMLST) phylogenetic tree of all available *E. coli* genomes recovered from Florida in Enterobase. CHeRI isolates from this study are identified in the outer ring (black). Source of isolation, or host, are identified in inner ring. Branch coloring is representative of *E. coli* phylogroup.

The minimum spanning tree of the greater Florida isolates showed that a majority of the CHeRI isolates were closely related to each other with a smaller subset of samples being phylogenetically dispersed (Figure 5, black). Within the B1 clade of the tree, isolates from three different ranches were identified to observe diversity patterns (Figure 5). Ranch 1 (purple) had single isolates from 7 different WTD – all of which had unique cgMLST genotypes (individual nodes on the tree). Ranch 2 (green) had 3 different deer, all with unique genotypes from each other. Within host, two WTD mortalities recovered 3 *E. coli* isolates from different organs. All within host isolates had the same genotype (Ranch 2 – Deer 2), however, the third deer (Ranch 2 – Deer 3) had two isolates with the same genotype and the third isolate was a separate phylogroup (A). In Ranch 3, similar mixed diversity patterns were observed within host, although the phylogroup was constrained to B1.

**Figure 5.**
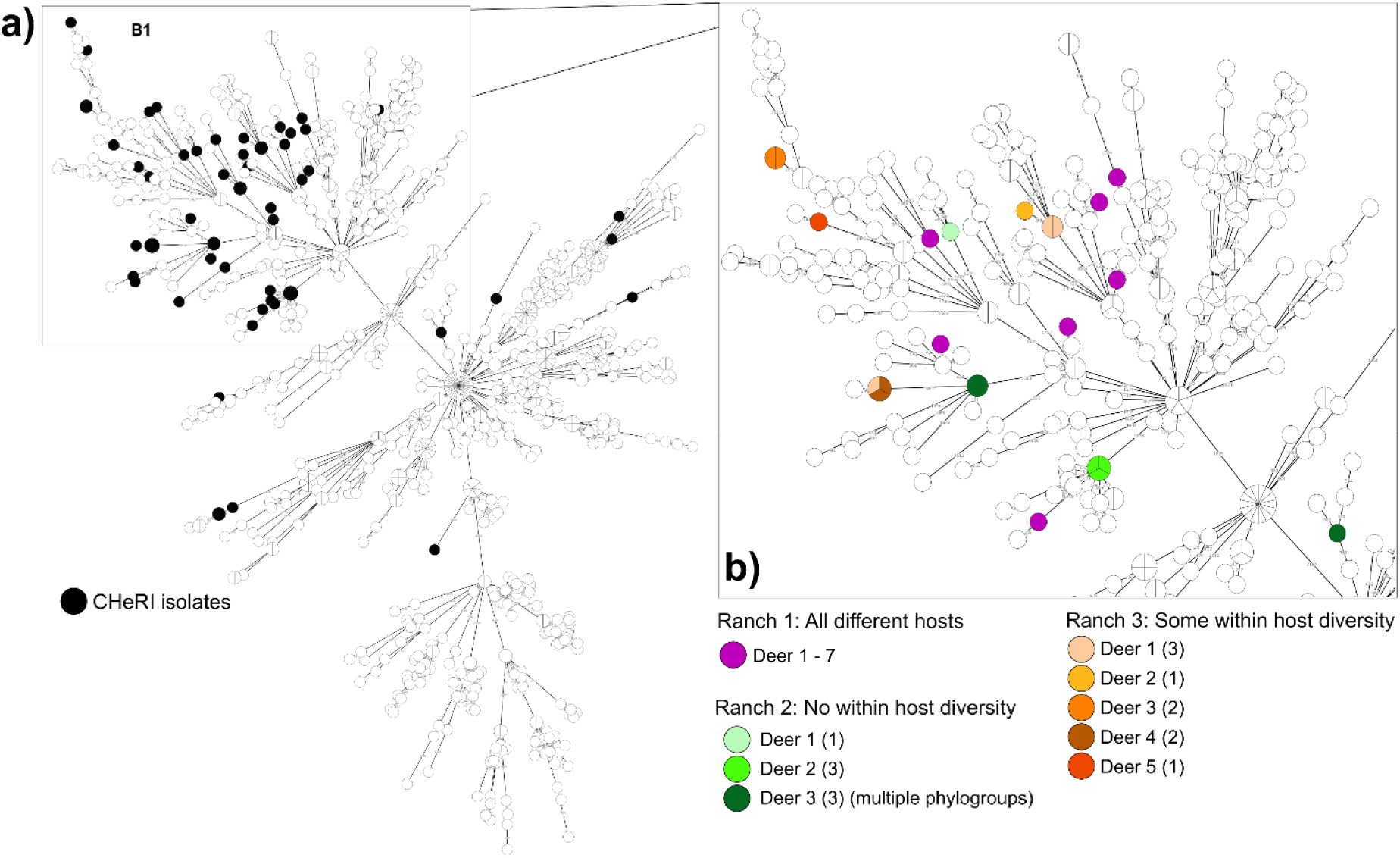
Minimum spanning tree of Florida *E. coli* genomes recovered from Enterobase. Location of CHeRI isolates within the greater Florida phylogeny (black nodes) (a). A subset of the tree, encompassing most of the B1 phylogroup branch, shows the phylogenetic distribution across three ranches in the 2023 study (b). Each ranch is an example of different within ranch and within host diversity scenarios.

## DISCUSSION

White-tailed deer ranches are distributed throughout most counties in Florida and participation in the University of Florida’s CHeRI project is widespread. However, there are noted gaps in participation (passive surveillance), particularly in southern and northeastern Florida. This northern region (Florida panhandle) has reported the greatest number of WTD mortalities throughout this project (2020 – 2022) – thus increased surveillance and a call for improved participation is suggested. Here, host co-infection by EHDV and/or BTV is not identified alongside a majority of the *E. coli* isolates. However, identification of these viral diseases, which typically act as primary infections [2], can be challenging if the infection is not active or recent. Situationally, this would occur if the host recovered from infection (low viral load) but was subsequently weakened and vulnerable to a secondary bacterial infection (commonly, systemic infections caused by *E. coli*). Often, *E. coli* strains are purely commensal, residing in the gut microbiome; however, intestinal pathogenic *E. coli* (InPEC) and extraintestinal pathogenic *E. coli* (ExPEC) strains can be major causes of human and animal disease [29,30]. Currently, only the Shiga toxin-producing *E. coli* (STEC) is a listed CDC reportable *E. coli* pathotype (CDC; National Notifiable Diseases Surveillance System (NNDSS) https://www.cdc.gov/nndss/index.html). However, additional virulence genes may be present in the *E. coli* genome that encode proteins to assist in host colonization and or subversion of the host immune system [31]. Many virulence genes arise in specific transmissible regions on the bacterial genome, called pathogenicity associated islands (PAIs) [32,33]. The combination of virulence genes produces phenotypes, termed pathotypes, defined as enteropathogenic (EPEC), enterohaemorrhagic (EHEC), or enterotoxigenic (ETEC) [32,34,35].

Here, we identified two strains with heat-stable endotoxin (ST) genes, classifying them as enterotoxigenic (ETEC) pathotypes. Additionally, one suspected *E. coli* isolate was later determined to be *Enterobacter hormaechei. Enterobacter hormaechei* is a Gram-negative bacterium that is primarily associated with clinical, or hospital-associated, infections [36,37]. This pathogen, and genetic near neighbors falling into the greater *Enterobacter cloacae* (ECC) complex, are causes of concern due to their high resistance to antimicrobials, specifically due to the production of β-lactamases [38]. Additionally, studies have identified resistance mechanisms in *Enterobacter* spp. that were also present in *E. coli* and some *Salmonella* spp. [37,39,40].

This study found that the most prevalent *E. coli* phylogroup amongst our WTD isolates was B1 (45/60 isolates). This phylogroup was also the most geographically widespread, with 17/20 ranches and 14/16 counties having at least one isolate classified as B1. Our second most prevalent phylogroup was A; in 7/60 isolates. Phylogroup B1 and A are less likely to cause disease (pathogenic) but can be associated with greater antimicrobial resistance (AMR) [8,10]. This is primarily resistance to ciprofloxacin and an increase in the production of β-lactamases – leading to resistance to penicillins, cephalosporins, cephamycins, monobactams and some carbapenems [41,42]. While a majority of isolates in this study fell into phylogroups B1 and A, a subset did fall into phylogroups B2 and D (4/60). These isolates were identified in 3/21 ranches, spanning 3/16 counties. Two of these counties, one located in central Florida and the other in the Florida panhandle, had isolates belonging to phylogroup B1. These findings highlight the risk for horizonal gene transfer of AMR genes, often identified in commensal strains (phylogroups A and B1), to more commonly pathogenic strains (phylogroups B2 and D) [10]. *E. coli* phylogroup distribution and prevalence in this study was comparable to a larger *E. coli* dataset retrieved from Enterobase, with no higher resolution geographic data.

Within the CHeRI project, previous studies targeting viral diseases EHDV and BTV found that seroprevalence of these viral diseases could be higher due to stocking of non-native mammalian species, potential amplifying hosts for EHDV, alongside white-tailed deer [43]. The mixed stocks of WTD and exotic ungulates, from both the Cervidae and Bovidae families, has been established practice on hunting preserves nationally [44]. Similarly, this co-mingling of different species and the ecological dynamics present within ranches may impact the exposures of WTD to *E. coli* species and phylogroups. Despite the zoonotic nature of this disease, there are limited studies specifically investigating *E. coli* populations, both pathogenic and commensal, in wild and domesticated ruminants. In a 2023 California study, Lagerstrom and Hadly identified a positive correlation between *E. coli* phylogroup diversity and range size in wild hosts [45]. This could not be significantly established in domesticated hosts due to limited data availability, however, these findings could suggest that domesticated hosts living on larger ranches, or ranches with more open ranging, may have opportunities for exposure to diverse *E. coli* populations [45]. Furthermore, the natural proximity of these domesticated hosts to human and built environments may lead to increased exposure to *E. coli* phylogroups more likely to impact or be found commensally in humans [45].

Despite the abundance of research and literature on *Escherichia coli*, the role of this bacterium in the public health domain can be challenging to define. Animal reservoirs play a major role in the transmission of pathogenic *E. coli* species into human hosts [46,47]. Often, cattle and dairy cows are linked to these cases [48,49], however, cervid hosts have also been suspected sources in human outbreaks. In 2011, an outbreak of human *E. coli* infections was traced back to strawberries likely contaminated with fecal material from black-tailed deer [50].

Co-mingling of WTD deer species with cattle has repeatedly led to the transmission of pathogenic *E. coli*, specifically O157 variants (Shiga toxin producing)(STEC) to WTD [46,51–53]. Understanding this disease in WTD populations can inform public health and safe hunting practice in Florida. Furthermore, elucidating the role of *E. coli*, or similar bacterial species, as secondary or primary sources of disease within these WTD populations is vital for maintaining the strength and breeding of these economically importance species across Florida ranches.

## Acknowledgements

This study was funded by the University of Florida, Institute of Food and Agricultural Science Cervidae Health Research Initiative (CHeRI), with funds provided by the State of Florida Legislature. We thank all of the participating ranches and CHeRI research personnel for sample collection and diagnostics.

## Notes

### Competing Interest Statement

The authors have declared no competing interest.

